# Cortico-Cerebellar Networks Drive Sensorimotor Learning in Speech

**DOI:** 10.1101/177527

**Authors:** Daniel R. Lametti, Harriet J. Smith, Phoebe Freidin, Kate E. Watkins

**Affiliations:** Department of Experimental Psychology The University of Oxford

**Keywords:** speech production, sensorimotor learning, tDCS, cerebellum, motor cortex

## Abstract

The motor cortex and cerebellum are thought to be critical for learning and maintaining motor behaviours. Here we use tDCS to test the role of the motor cortex and cerebellum in sensorimotor learning in speech. During productions of ‘head’, ‘bed’, and ‘dead’, the first formant of the vowel sound was altered in real-time towards the first formant of the vowel sound in ‘had’, ‘bad’, and ‘dad’. Compensatory changes in first and second formant production were used as a measure of motor adaptation. TDCS to either the motor cortex or the cerebellum improved sensorimotor learning in speech compared to sham stimulation. However, in the case of cerebellar tDCS, production changes were restricted to the source of the acoustical error (i.e. the first formant). Motor cortex tDCS drove production changes that offset errors in the first formant, but, unlike cerebellar tDCS, adaptive changes in the second formant also occurred. The results suggest that motor cortex and cerebellar tDCS have both shared and dissociable effects on motor adaptation. The study provides initial causal evidence in speech production that the motor cortex and the cerebellum support different aspects of sensorimotor learning. We propose that motor cortex tDCS drives sensorimotor learning towards previously learned patterns of movement, while cerebellar tDCS focuses sensorimotor learning on error correction.

## Introduction

Speech might be the most complicated human behaviour. In one second of fluid speech, we produce 2 to 3 words made up of 10 to 12 phonemes (Levelt, 1999). This task demands the close coordination of at least a dozen muscles, and, simultaneously, careful monitoring of auditory and somatosensory feedback to ensure that speech goals are achieved (Houde and Nagarajan, 2011; Tourville and Guenther, 2011). To date, the neural basis of motor learning has been underexplored in speech compared to other motor behaviours such as reaching. The motor cortex and cerebellum are thought to be critical for the learning and maintenance of movements. Sensorimotor learning alters networks within both structures (Albert et al., 2009), and the neural excitability of the cerebellum tracks adaptive changes in movement (Jayaram et al., 2011; Schlerf et al., 2012a). Here, for the first time, we use transcranial direct current stimulation (tDCS) to test the role of the motor cortex and cerebellum in sensorimotor learning during speech production.

Over the last twenty years, sensorimotor learning has been extensively studied by having human participants interact with manipulanda such as robotic arms, styli, and joysticks. In a typical study, participants use one of these devices to control a cursor on a computer screen. After a series of movements to targets arranged around a circle, a consistent rotation of the cursor’s position is introduced and participants must adapt their movements to reach the targets (Krakauer et al., 2000). This paradigm, which is a form of visuomotor adaptation akin to mirror drawing, has become a standard experimental model for investigating sensorimotor learning. However, a parallel line of work has recently developed an experimental model for studying sensorimotor learning in speech (Houde and Jordan, 1998; Purcell and Munhall, 2006; Rochet-Capellan and Ostry, 2011).

Sensorimotor learning in speech is studied by altering vowel sounds in real-time as speech is produced. Vowel sounds are defined by peaks in their acoustical spectra, in particular the relationship between the first and second formants (Ladefoged, 1975). In a typical experiment, participants produce consonant-vowel-consonant words into a microphone and their speech is played back to them over headphones with an imperceptible delay. To induce sensorimotor learning, acoustical effects processors are used to alter the formant structure of the vowel (Houde and Jordan, 1998). For instance, the first formant may be increased or decreased so that participants say a word like “head” but hear themselves producing something that sounds closer to “had” or “hid” (Rochet-Capellan and Ostry, 2011; Lametti et al., 2014a). When such an alteration is experienced, participants learn to adapt their speech production to compensate for the perceived error. This type of sensorimotor learning produces reliable patterns of adaptation and after-effects—and, as an experimental model of motor learning, it has important advantages over visuomotor adaptation.

Sensorimotor learning in speech is driven by alterations in a participant’s’ own voice, as opposed to alterations in an abstract representation of limb position (e.g. a cursor) via an interaction with an external device (e.g. a joystick). Participants are seldom aware of altered auditory feedback and learning is not affected by explicit movement strategies. For example, participants who are instructed to avoid compensating for altered auditory feedback, or who have full knowledge of the manipulation, show the same patterns of learning as naive participants (Munhall et al., 2009). This suggests that sensorimotor learning in speech is largely—if not entirely—driven by implicit changes in motor systems, as opposed to the combination of implicit changes and explicit cognitive strategies that shape visuomotor learning (Mazzoni and Krakauer, 2006; Benson et al., 2011; Taylor et al., 2014; McDougle et al., 2016).

Another advantage of studying sensorimotor in speech is that more complex movements and corresponding patterns of adaptation can be examined. Studies of visuomotor adaptation in limb movement typically constrain movements to two joints, whereas sensorimotor learning in speech is unconstrained. Compensation for altered auditory feedback can be accompanied by changes in production that seem unrelated to the induced acoustical error. Alterations in the first formant (F1) of vowel sounds lead to compensation in F1, but they frequently cause changes in the second formant (F2) as well (MacDonald et al., 2011; Rochet-Capellan and Ostry, 2011). In this case, changes in F2 are not related to biomechanical constraints—that is, F1 and F2 can be altered independently (Villacorta et al., 2007; MacDonald et al., 2011). Here we posit that the F2 changes that occur following F1 alterations may serve to place articulation closer to a previously learned speech movement.

One disadvantage of studying sensorimotor learning in speech is that patterns of compensation for altered auditory feedback are more variable than compensation patterns observed during visuomotor learning. In a typical altered feedback experiment, 10-40% of participants fail to show learning, and those that do learn rarely eliminate more than 30% of the experimentally induced acoustical error (Lametti et al., 2012, 2014a; Rochet-Capellan et al., 2012). This added variability means that more participants must be tested—but it eliminates the possibility that effects are missed because learning rapidly reaches completion. It may also better reflect the high degree of variability observed in real-world motor learning tasks.

The brain-basis of sensorimotor learning in speech has been understudied compared to sensorimotor learning associated with reaching movements. Functional neuroimaging suggests that novel speech production is associated with increases in activity in both traditional motor areas (e.g. frontal lobe motor areas and the basal ganglia) as well as areas implicated in the feedback-based maintenance of motor behaviours (e.g. premotor cortex and the cerebellum) (Rauschecker et al., 2008; Segawa et al., 2015). Patients with impaired cerebellar function show deficits in adaptation related to predictable alterations of auditory feedback (Parrell et al., 2017). Causal interventions involving transcranial magnetic and direct current stimulation have recently linked predictive language processing to the cerebellum (Lesage et al., 2012; Miall et al., 2016). However, causal investigations of the brain-basis of sensorimotor learning in speech remain limited in healthy participants (Shum et al., 2011).

A significant number of studies have explored the impact of single-session tDCS on visuomotor adaptation. In many cases, the motor cortex, the cerebellum, or both are targeted (Galea et al., 2011; Panouillères et al., 2015). Typically, 1 to 2 mA of direct current is applied to the brain for 15 to 20 minutes. Anodal stimulation is frequently used and it is applied online—that is, the anode is placed over the brain area of interest and stimulation is delivered throughout the adaptation task. The precise mechanisms by which tDCS might alter the brain-basis of sensorimotor adaptation are not completely understood. TDCS is thought to change the excitability of neurons in the stimulated brain area, although effects at the network level are also observed (Bestmann et al., 2015).

Demonstrating a reliable impact of either cerebellar or motor cortex tDCS on an aspect of sensorimotor learning (e.g. the amount of learning) is important for at least three reasons: consistent effects would 1) provide insight into the role of these brain areas in motor behaviour; 2) bolster the still-disputed idea that stimulating the brain with a weak electric current (e.g. tDCS) is a reliable tool for investigating brain-behaviour relationships (Horvath et al., 2015); and 3) have important clinical implications related to motor rehabilitation (Allman et al., 2016). Unfortunately, the results of studies examining the impact of anodal tDCS on visuomotor adaptation have been inconsistent. Initial work suggested that anodal tDCS to the cerebellum speeds up visuomotor learning (Galea et al., 2011), but recent studies have failed to consistently replicate this result (Jalali et al., 2017). Others have noted improvements in visuomotor adaptation when learning is paired with motor cortex tDCS (Panouillères et al., 2015), but, again, this result has not been consistently observed (Doppelmayr et al., 2016).

For the reasons noted above, we argue that adaptation to altered auditory feedback presents a more ecologically valid experimental model of sensorimotor learning. To advance our understanding of the neural basis of speech production in healthy participants, and help address the mixed results in the literature related to the impact of tDCS on sensorimotor learning, we applied anodal tDCS to either the motor cortex or the cerebellum during sensorimotor learning in speech. We examined the amount and the maintenance of learning following altered auditory feedback between stimulation locations, and compared to a group of participants who received sham stimulation to both brain areas.

There are known reciprocal links between the cerebellum and primary motor areas (Orioli and Strick, 1989; Kelly and Strick, 2003). Indeed, both tDCS and sensorimotor learning cause changes in cerebellar excitability that are reflected in the strength of the cerebellum’s influence on the excitability of primary motor areas (Galea et al., 2009; Jayaram et al., 2011; Schlerf et al., 2012a). A possibility, then, is that tDCS to *either* structure impacts cortico-cerebellar networks involved in updating the neural commands that control movements. Given the mixed results in the literature and the known reciprocal links between the two simulated structures, we ran the experiment with the simple hypothesis that tDCS would improve sensorimotor learning in speech. That is, we did not predict that stimulating a particular point of the cortico-cerebellar circuit (i.e. the motor cortex or the cerebellum) would cause improvements in learning over stimulation of any other point.

## Methods

### Participants and Apparatus

Sixty participants between the ages of 18 and 35 years were recruited from the Oxford University community. Twenty-six of the participants were male. Participation was restricted to right-handed fluent English speakers with normal hearing and speech. The Central University Research Ethics Committee approved the experimental protocol. Prior to testing, participants gave their informed consent and were screened for criteria that would exclude them from tDCS.

The altered auditory feedback system used in the study was similar to that depicted in Figure 1 of Rochet-Capellan and Ostry (2011). Participants wore over-ear headphones (Sennheiser) and produced words that appeared on a computer screen into a head-mounted microphone (Shure). An acoustical effects processor (TC Helicon) and analog filters (Rockland) were used to alter speech and play it back to participants with a minimal delay. A direct current stimulator (NeuroConn) applied stimulation to either the cerebellum or motor cortex.

**Figure 1:**
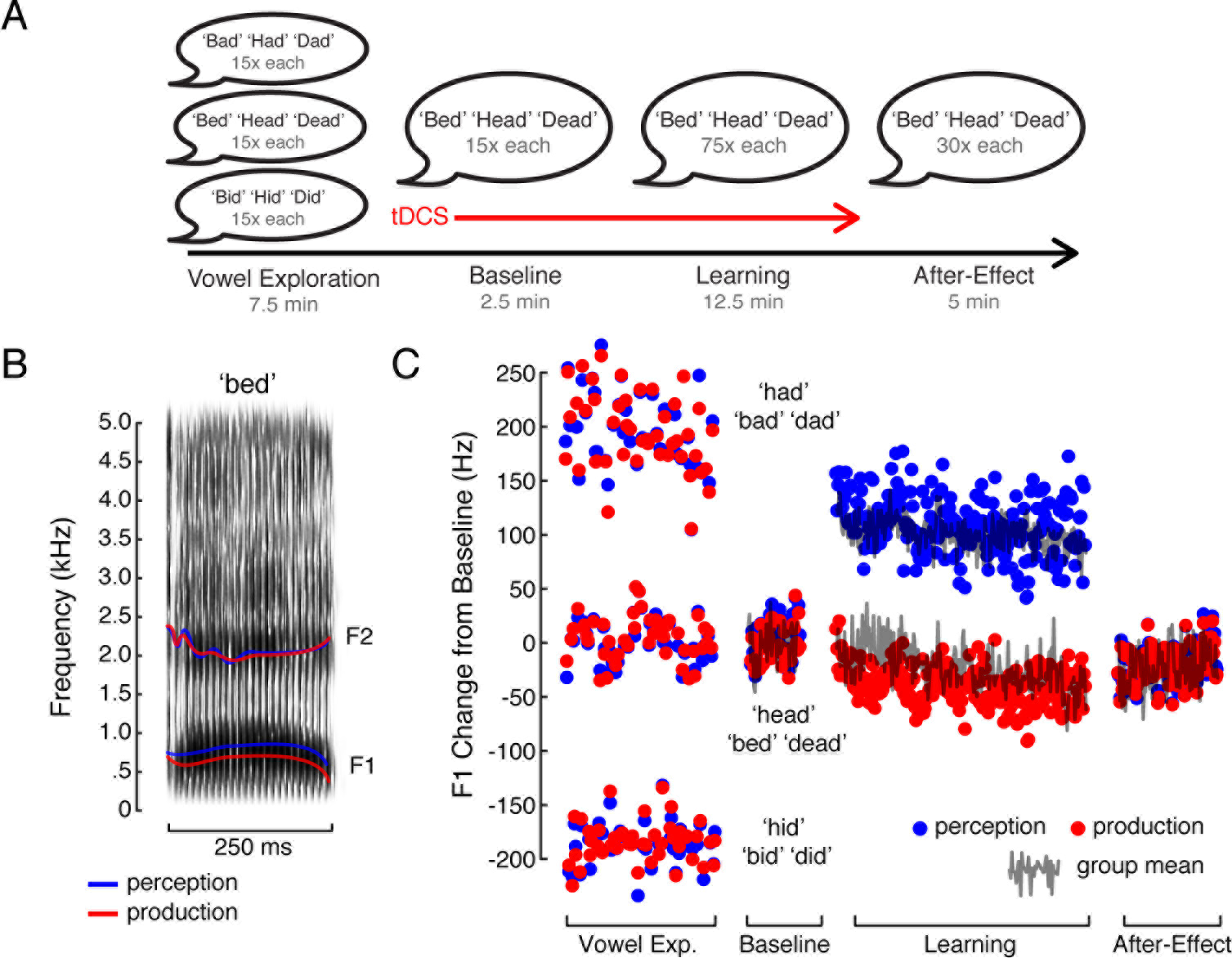
Experimental design (A) The study had four phases: Vowel Exploration, Baseline, Learning and After-Effect in which participants produced words as they appeared on a screen. Anodal tDCS was applied during Baseline and Learning. (B) The spectrogram depicts the formant structure for an utterance produced in the study. The first and second formants are highlighted by the coloured lines. The red line shows the first and second formant that the participants produced, whereas the blue lines show the first and the second formant that the participant heard during altered auditory feedback. (C) Change in first formant (F1) frequencies relative to baseline F1 frequency for a single participant. The blue circles represent the F1 the participant heard and the red circles represent the F1 the participant produced. For comparison purposes, the solid grey line represents the change in F1 production over the Baseline, Learning, and After Effect phases of the experiment for the group that received cerebellar stimulation.

### Procedure

The experiment was divided into blocks of utterances: Vowel Exploration, Baseline, Learning, and After-Effects (Figure 1A and 1C). Within each block, consonant-vowel-consonant words were presented on a computer screen for 1500 ms, one at a time. Words were presented in a pseudorandom order, such that each word was produced an equal number of times. The inter-trial-interval was 750 ms. Participants were instructed to produce the words as they appeared on the screen so that they could hear themselves through the headphones.

The experiment began with unaltered speech. Participants produced ‘dead’, ‘bed’, ‘head’, ‘dad’, ‘bad’, ‘had’, ‘did’, ‘bid’, and ‘hid’ 15 times each. This first block of utterances (Vowel Exploration) was designed to let participants explore the vowel space they would later experience altered feedback in. The stimulating electrodes were then attached to the head and the direct current stimulator was turned on. Following a 60-s break in which the current ramped up (30-s) and participants confirmed that the stimulation was comfortable (30-s), the words ‘head’, ‘bed’, and ‘dead’ were produced 15 times each. This block of utterances (Baseline) was designed to give a measure of speech production with stimulation, prior to sensorimotor learning. Participants then produced ‘head’, ‘bed’, and ‘dead’ 75 times each with altered auditory feedback (Learning). The experiment concluded with unaltered productions of ‘head’, ‘bed’, and ‘dead’ in which each word was produced 30 times. This final block of production (After-Effect) was designed to provide a measure of the maintenance of sensorimotor learning in speech after altered feedback was removed. With the exception of the 60-s break between the first and second baseline blocks, there were 30-s breaks every 45 utterances.

### Sensorimotor Learning

To induce sensorimotor learning in speech, the first formant frequency (F1) of the vowel sound in the words ‘head’, ‘bed’, and ‘dead’ was increased by 21% (4.0% SD) (Figure 1B and 1C). Altered F1 productions were mixed with the rest of the unaltered speech signal and 60-dB speech-shaped masking noise and then played back to participants through the headphones with an unnoticeable delay. The effect of the manipulation was that participants produced ‘head’, ‘bed’, and ‘dead’ into the microphone but heard themselves (via the headphones) producing words that sounded more like ‘had’, ‘bad’, and ‘dad’. Participants adapt to this sensory-motor mismatch by changing their formant productions to correct the perceived error (e.g. Houde and Jordan, 1998; Purcell and Munhall, 2006; Munhall et al., 2009; Rochet-Capellan and Ostry, 2011; Lametti et al., 2012, 2014a, 2014b; Shiller and Rochon, 2014).

### Direct Current Stimulation

Transcranial direct current stimulation (tDCS) was applied to either the right cerebellum or left hemisphere motor areas during the Baseline and Learning phases of the experiment. A 2-mA current was delivered through 5 by 7 cm electrodes placed within saline-soaked sponges. The sponges were positioned on the scalp using flexible self-adherent bandage (Cobane, 3M) prior to the start of the Baseline phase. One bandage ran from below the inion around the forehead and back to below the inion. A second bandage ran from the bottom of the jaw around the top of the head (in front of the vertex) and back to the bottom of the jaw. This setup was used in all conditions. The electrodes were positioned under the bandages such that, in the case of motor cortex stimulation, with the electrode in a portrait configuration, the top left corner of the anode was positioned at a point one third of the length along a line connecting the vertex to the tragus, placing it over articulator representations in primary motor cortex. The cathode was placed in landscape configuration on the forehead above the right eye. In the case of cerebellar stimulation, with the electrode in a portrait configuration, the center of the anode was positioned 3 cm lateral to the inion on the right side of the scalp and the cathode was placed in landscape configuration on the right cheek (Galea et al., 2011; Lametti et al., 2016).

Thirty participants received stimulation to the cerebellum and thirty participants received stimulation to the motor cortex. In each group, 20 of the 30 participants received 16 minutes of real (or effective) stimulation and the remaining 10 received 90 seconds of sham (or ineffective) stimulation. In both cases, the current was ramped up to 2 mA over 30-s and down to zero over 30-s. Put another way, during sensorimotor learning, 20 participants received real motor cortex stimulation (Motor Cortex group), 20 participants received real cerebellar stimulation (Cerebellar group), and 20 participants received sham stimulation (Sham group), with an equal divide in the Sham group between a cerebellar and motor cortex electrode arrangement. The participants and the researcher were blind to the stimulation condition. A third party, who was not involved in the experimental setup, randomly assigned participants to each group and operated the stimulator.

### Auditory Analysis

The formant analysis closely follows previous work (Bourguignon et al., 2014; Lametti et al., 2014a; Shiller and Rochon, 2014). Speech was recorded at 22050 Hz. A 50-ms segment at the centre of each vowel was extracted. The mean first and second formant frequencies across this segment were calculated using Linear Predictive Coding (LPC) analysis (Figure 1B). LPC parameters were selected on a per-subject basis to minimize variance in F1 and F2. First and second formant values greater than 3 standard deviations from a participant’s mean F1 and F2 values were excluded (< 0.6% of the data). Formant extraction and data analysis was performed in Matlab (Mathworks).

The three groups of participants had acoustically similar speech prior to sensorimotor learning. Baseline F1 frequency for the words ‘head’, ‘bed’, and ‘dead’ averaged 701 Hz for the Motor Cortex group, 690 Hz for the Cerebellar group, and 681 Hz for the Sham group (ANOVA comparing baseline F1: F(2, 57) = .224, p = .80). Baseline F2 frequency for the words ‘head’, ‘bed’, and ‘dead’ averaged 1810 Hz for the Motor Cortex group, 1893 Hz for the Cerebellar group, and 1815 Hz for the Sham group (ANOVA comparing baseline F2: F(2, 57) = 1.53, p = .23). There were no significant correlations between baseline F1 and F2 frequency and the amount of learning or learning-related after-effects in each formant (r < 0.2 in all cases).

### Planned Analyses

Sensorimotor learning was quantified on a per-subject basis as the change in produced F1 and F2 formant frequencies from the baseline phase of the experiment in response to altered feedback (Houde and Jordan, 1998; Rochet-Capellan and Ostry, 2011; Bourguignon et al., 2014; Lametti et al., 2014a, 2014b). Specifically, the average change from baseline in produced F1 and F2 during the last 30 utterances of altered auditory feedback and the first 15 utterances of the after-effect phases of the experiment were calculated and used as measures of sensorimotor learning (respectively). Within groups, changes in F1 and F2 frequency associated with learning were assessed using t-tests. In this case, directional t-tests were used, as the direction of formant frequency change that would constitute sensorimotor learning in response to a perceived F1 increase was predictable. Between groups, changes in F1 and F2 were compared using repeated measures ANOVA and, within the two phases of the experiment (learning and after-effects), one-way ANOVA. The data were checked for normality using the Shapiro-Wilk test and found to be normally distributed. Post-hoc comparisons were performed using two-tailed t-tests and corrected for multiple comparisons using the Bonferroni-Holm method. The significance level for all statistical tests was p < 0.05.

### Unplanned Analyses

Sensorimotor learning in speech in response to F1 alterations is often accompanied by learned changes in F2. Learning involves both precise corrections for acoustical errors and more general changes that may steer production towards previously learned motor plans in the F1-F2 vowel space. We explored this idea in the data. Specifically, the impact of sensorimotor learning and tDCS on the distance in F1-F2 space between ‘head’, ‘bed’, and ‘dead’ productions and ‘hid’, ‘bid’, and ‘did’ productions was examined. The aim of this analysis (which was planned based on the results in Figures 2 and 3) was to test the degree to which tDCS and altered auditory feedback drove a change in participants’ productions towards F1 and F2 values associated with ‘hid’, ‘bid’, and ‘did’. To do this, in F1 and F2 space, we calculated the Euclidean distance between participants’ productions of ‘head’, ‘bed’, and ‘dead’ during the main experiment and their productions of ‘hid’, ‘bid’, and ‘did’ during the vowel exploration phase. Learning- and after-effect-related changes in this distance from baseline productions of ‘head’, ‘bed’, and ‘dead’ were compared between groups using repeated measures ANOVA and one-way ANOVA. Within the learning and after-effect phases of the experiment, post-hoc comparisons were performed using two-tailed t-tests, and corrected for multiple comparisons using the Bonferroni-Holm method.

### Data Availability

The individual data points used in all statistical tests are shown in the manuscript (see Figures 2B, 3B, and 4C). The raw formant data is available upon request.

**Figure 2:**
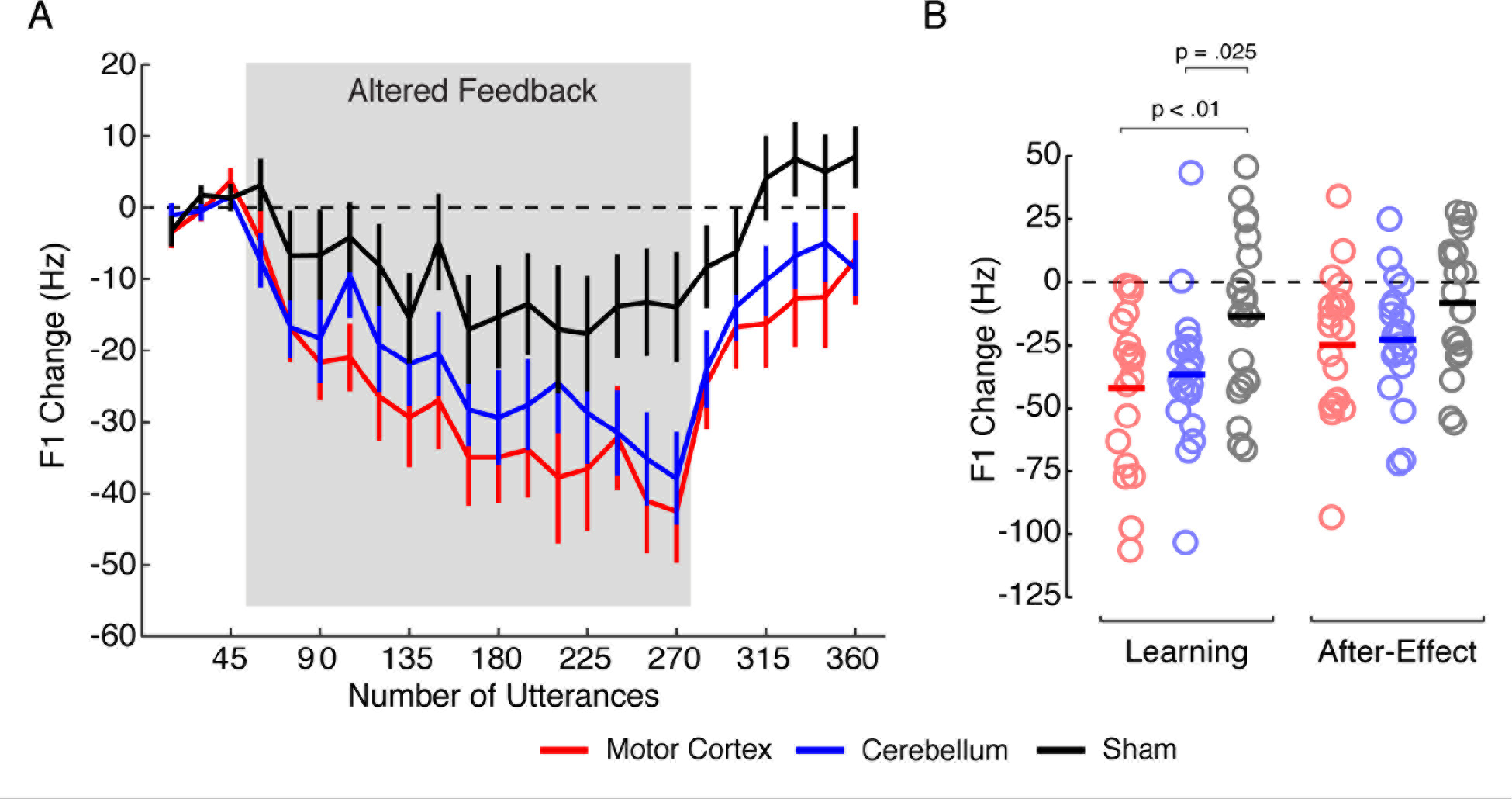
Changes in F1 production related to sensorimotor learning and tDCS location (A) Changes in produced F1 frequency in response to altered F1 feedback are shown for the baseline, sensorimotor learning and after-effect phases of the experiment. Each data point represents the mean of 15 utterances of ‘bed’ ‘dead’ and ‘head’. Error bars show +/− a standard error. The dashed line reflects no change from baseline. (B) Mean changes in F1 frequency are shown for the end of the learning phase (utterances 241 to 270) and the start of the after-effect phase (utterances 271 to 285). The solid bars represent group means and the open circles represent data from individual participants. Significant between-groups differences are indicated with p-values. In both panels, red, blue and black lines/circles represent the motor cortex, cerebellar, and sham tDCS groups.

## Results

The purpose of this study was to investigate the role of the cerebellum and motor cortex in sensorimotor learning in speech. Two groups of twenty participants each received anodal transcranial direct current stimulation (tDCS) to either the motor cortex or the cerebellum during a speech adaptation task in which the first formant frequency (F1) of their ‘head’, ‘bed’, and ‘dead’ productions was increased in real-time. The extent of compensatory changes in formant frequency production, and the maintenance of these changes (after-effects) following removal of altered auditory feedback were compared between the two groups, and with a group of twenty different participants who received sham tDCS during the same task.

Figure 2A shows how produced F1 frequency changed across the sensorimotor learning (utterances 46 to 270) and after-effect (utterances 271 to 360) phases of the experiment, in comparison with baseline measures of F1 frequency. The red, blue and black lines represent production changes associated with receiving motor cortex, cerebellar or sham tDCS, respectively. Change-in-F1 frequency at the end of the learning phase was significantly different from baseline. On average, participants in each group altered the frequency of their F1 productions in a downward direction to compensate for the perceived upward shift in F1 (p < .05, in each case). This indicates that adaptation to the feedback alteration occurred in all stimulation conditions. Adaptation in the motor cortex, cerebellar, and sham groups offset an average of 29%, 26% and 10% of the F1 error (respectively).

Transcranial direct current stimulation (tDCS) drove a difference in the extent of F1 adaptation between the stimulation conditions (F(2,57) = 3.87, p = .027; main effect of group i.e. stimulation condition). The interaction between experiment phase and stimulation condition did not reach statistical significance. That is, the pattern of F1 adaptation between the groups at the end of learning was largely mirrored in the after-effect trials. Figure 2B shows changes in F1 production at the end of the sensorimotor learning phase of the experiment (left panel) and at the start of the after-effect phase (right panel). The open circles represent data from individual participants during these two phases. As in Figure 2A, participants receiving motor cortex, cerebellar and sham tDCS are shown in red, blue and black, respectively. By the end of altered auditory feedback there were differences in the amount of learning between the groups (F(2,57) = 4.65, p = .013). Change-in-F1 frequency from baseline was greater in the cerebellar tDCS group compared with the sham group (t(38) = −2.34, p = .025), and in the motor cortex tDCS group compared with the sham group (t(38) = −2.76, p < .01). However, change-in-F1 frequency was not significantly different between the cerebellar and motor cortex groups (t(38) = .56, p = .59). This pattern of results appeared similar in the after-effect phase of the experiment but between-group differences were not statistical significance (F(2,57) = 2.29, p = .11). Taken together, these results suggest that learned changes in the frequency of F1 productions in response to F1 alterations were improved by tDCS regardless of whether the cerebellum or the motor cortex is stimulated.

Sensorimotor learning associated with F1 alterations can drive learned changes in both the first and second formants. Previous work has noted that upward manipulations of F1 frequency lead to compensatory *decreases* in produced F1 frequency and *increases* in produced F2 frequency (MacDonald et al., 2011; Rochet-Capellan and Ostry, 2011). We thus looked for increases in F2 frequency associated with our F1 manipulation.

Figure 3A shows how produced F2 frequency changed across the sensorimotor learning (utterances 46 to 270) and after-effect (271 to 360) phases of the experiment in response to an alteration in F1. By the end of sensorimotor learning, there was a difference in the extent of F2 frequency change between the three stimulation conditions (F(2,57) = 4.72, p = .013; main effect of group i.e. stimulation condition. The interaction between experiment phase and stimulation condition was not significant.) The groups that received motor cortex and sham stimulation showed an increase in produced F2 frequency (compared with baseline) over the sensorimotor learning and after-effect phases of the experiment (p < .05, in each case). An increase in F2 frequency was not observed in the group that received cerebellar stimulation (p = 0.2).

**Figure 3:**
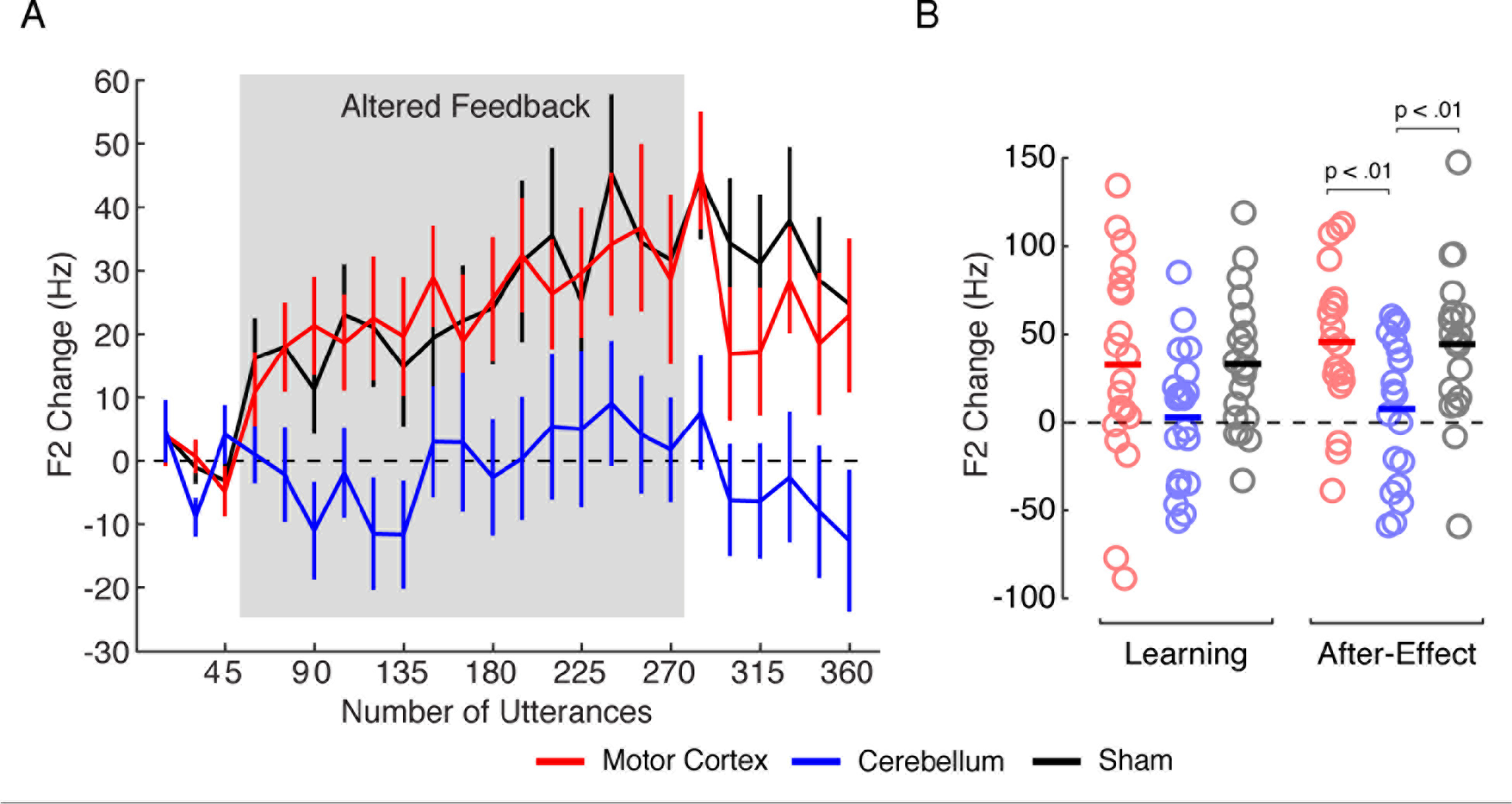
Changes in F2 production related to sensorimotor learning and tDCS location (A) Changes in produced F2 frequency in response to altered F1 feedback are shown for the baseline, sensorimotor learning, and after-effect phases of the experiment. Each data point represents the mean of 15 utterances of ‘bed’ ‘dead’ and ‘head’. Error bars show +/− a standard error. The dashed line reflects no change from baseline. (B) Mean changes in F2 frequency are shown for the end of the learning phase (utterances 241 to 270) and the start of the after-effect phase (utterances 271 to 285). The solid bars represent group means and the open circles represent data from individual participants. Significant between-groups differences are indicated with p-values. In both panels, red, blue and black lines/circles represent the motor cortex, cerebellar, and sham tDCS groups.

Figure 3B shows changes in produced F2 frequency at the end of the sensorimotor learning phase of the experiment (left panel) and at the start of the after-effect phase of the experiment (right panel), in response to the F1 alteration. As in Figure 2B, the open circles represent data for individual participants at these phases of the experiment. By the end of learning, the motor cortex and sham groups appeared to show a greater increase in F2 frequency than the cerebellar group but these differences did not reach statistical significance (F(2,57) = 2.79, p = .070). Group differences were statistically significant in the after-effect phase of the experiment (F(2,57) = 5.70, p < .01). Specifically, the motor cortex and sham groups showed a reliably greater increase in F2 frequency compared to the cerebellar group (t(38) = 2.94, p < .01 and t(38) = 2.79, p < .01, respectively). These results suggest that cerebellar tDCS inhibited a change in produced F2 frequency normally associated with adaptation to alterations in F1 frequency.

Taken together, the findings in Figure 2 and 3 indicate that cerebellar and motor cortex tDCS have both shared and dissociable effects on sensorimotor learning in speech. Cerebellar stimulation restricted learned changes in speech production to the source of the acoustical error, in this case F1. In contrast, motor cortex stimulation lead to learned changes in both F1 and F2 production. Why might an experimentally induced change in F1 drive a learned change in both F1 and F2? One possibility is that sensorimotor learning in speech involves both precise corrections for acoustical errors and more general changes intended to steer productions towards previously learned motor plans. Figure 4 examines this idea in relation to motor cortex and cerebellar tDCS. Specifically, via a post-hoc analysis, we explored the degree to which patterns of compensation between the stimulation conditions moved participants’ productions towards previous produced speech utterances in the F1-F2 vowel space.

**Figure 4:**
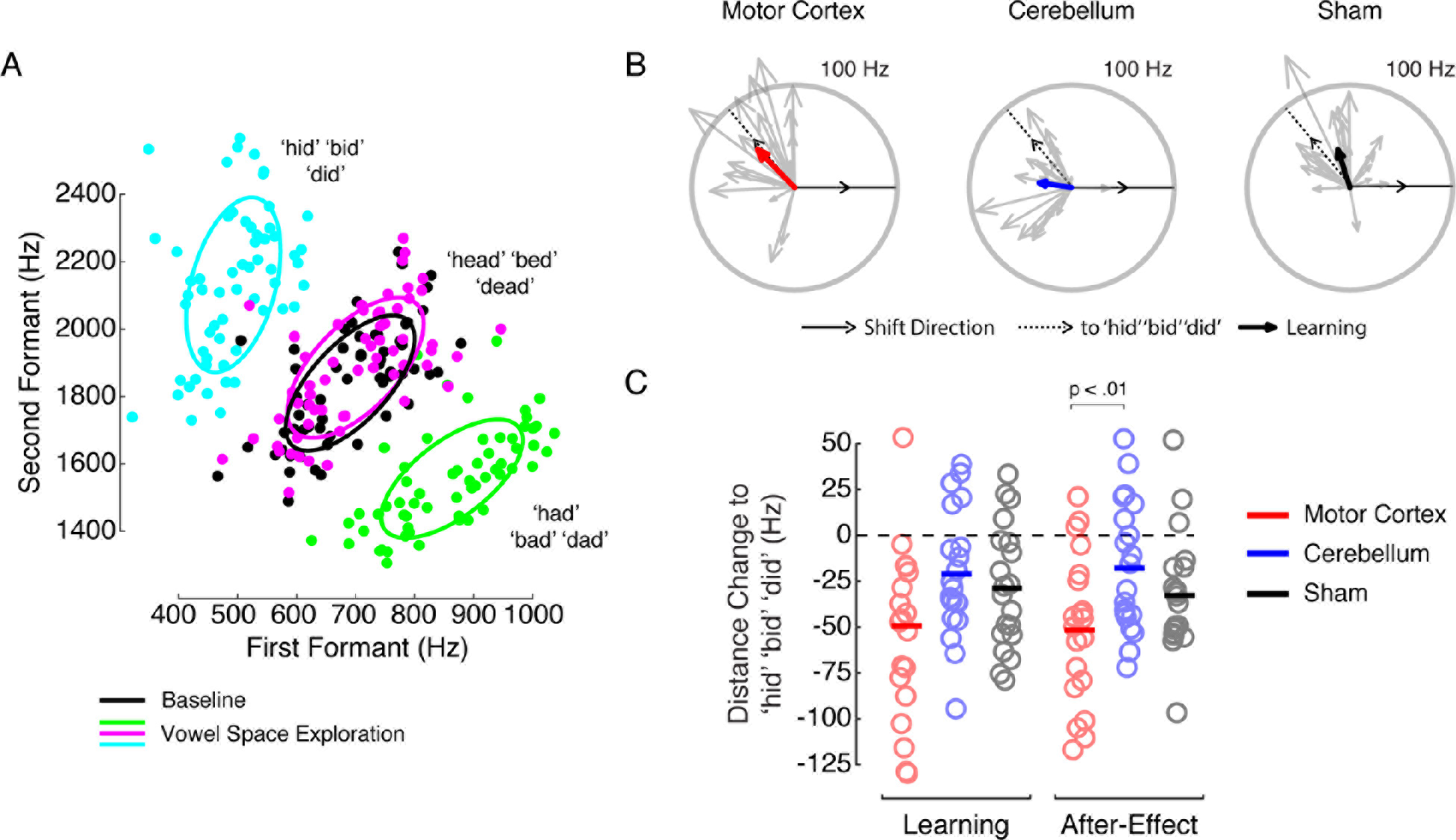
(A) Mean F1 and F2 values for ‘hid’ ‘bid’ ‘did’ (cyan), ‘head’ ‘bed’ ‘dead’ (magenta/black) and ‘had’ ‘bad’ ‘dad’ (green) from the Vowel Exploration and Baseline phase of the study. The ellipses represent a 50% confidence interval around the data points of similar colour. (B) Direction and magnitude of learned production changes in F1-F2 space for each stimulation condition. The grey arrows show the average production change over the end of the learning and after effect phases of the experiments for individual participants. The coloured arrows show the average production change across the group. The thin black line shows the direction of the induced F1 alteration. The dashed line reflects the direction of production change that would move productions from ‘head’ ‘bed’ ‘dead’ towards ‘hid’ ‘bid’ ‘did’ in F1-F2 space. (C) The change in Euclidean distance in Hz (in F1-F2 space) between ‘head’ ‘bed’ ‘dead’ and ‘hid’ ‘bid’ ‘did’ in relation to learning and brain stimulation. Negative values indicate that production has moved closer to ‘hid’ ‘bid’ ‘did’. The left side of the figure shows this change at the end of learning; the right side of the figure shows learning-related after-effects. The solid bars represent group means and the open circles represent data from individual participants. Significant between-groups differences are indicated with p-values. The dashed line represents no change in the euclidean distance from baseline. Red, blue and black circles represent the motor cortex, cerebellar, and sham tDCS groups.

Figure 4A shows patterns of F1 and F2 production during the vowel exploration and baseline phases of the experiment. The filled circles represent the mean frequency of F1 and F2 productions associated with the words ‘hid’ ‘bid’ ‘did’, ‘head’ ‘bed’ ‘dead’, and ‘had’ ‘bad’ ‘dad’, for each of the 60 participants. To induce sensorimotor learning, baseline F1 frequency associated with the words ‘head’ ‘bed’ and ‘dead’ (black circles) was altered towards an F1 frequency associated with ‘had’ ‘bad’ ‘dad’ (green circles). In response, participants learned to decrease the frequency of their F1 productions. Compared to strict F1-only compensation, an increase in F2 concurrent with this learned decrease in F1 would place productions closer to the ‘hid’ ‘bid’ ‘did’ region of the vowel space.

Figure 4B shows how the first and second formant frequency of ‘head’ ‘bed’ and ‘dead’ productions changed in the F1-F2 vowel space, in relation to the F1 alteration and brain stimulation. The vectors represent the direction and magnitude of the compensatory change from baseline. The grey vectors represent data from individual participants and the coloured vectors represent the average vector for each stimulation condition. The dashed lines represent the direction in F1-F2 space in which productions would have to change to move speech from the ‘head’ ‘bed’ ‘dead’ region of the vowel space to the ‘hid’ ‘bid’ ‘did’ region of the vowel space.

Sensorimotor learning in the motor cortex (red arrow) stimulation condition appeared to move participants’ productions closer to ‘hid’ ‘bid’ ‘did’ than learning in the cerebellar (blue arrow) stimulation condition. This observation is quantified in Figure 4C. The figure shows the change in Euclidean distance between ‘head’ ‘bed’ ‘dead’ and ‘hid’ ‘bid’ ‘did’ in F1-F2 space. Learning-related changes in this distance differed between the three stimulation conditions (F(2,57) = 3.61, p = .033; main effect of group i.e. stimulation condition. The interaction between experiment phase and stimulation condition was not statistically significant.) By the end of learning, the frequency of F1 and F2 productions in the motor cortex group was 49.5 Hz closer in F1-F2 space to the frequency of F1 and F2 productions associated with ‘hid’ ‘bid’ ‘did’. Between group differences in this measure reached statistical significance in the after-effect phase of the experiment (F(2,57) = 4.35, p = .017). Specifically, in the case of the group that received motor cortex stimulation, the reduction in distance between F1 and F2 frequencies associated with ‘head’ ‘bed’ ‘dead’, and F1 and F2 frequencies associated with ‘hid’ ‘bid’ ‘did’, was greater than that observed in the cerebellar group (t(38) = 2.80, p < .01). This analysis provides preliminary evidence that, compared to cerebellar stimulation, motor cortex stimulation drove sensorimotor learning in speech closer to a known pattern of speech production.

## Discussion

We examined the impact of tDCS on sensorimotor learning in speech production. Participants produced the words ‘head’, ‘bed’, and ‘dead’ into a microphone, and the first formant frequency of the vowel sound was increased and played back via headphones with an unnoticeable delay. Changes in production associated with this manipulation were compared between three groups of twenty different participants who received anodal tDCS during the task. In two groups, sixteen minutes of 2.0 mA tDCS was delivered to the motor cortex or the cerebellum; a third group received sham stimulation to either the motor cortex or the cerebellum. In our sample, stimulation to both sites drove increases in F1 compensation compared to sham stimulation. Cerebellar tDCS restricted compensation to the source of the acoustical error (i.e. the first formant), whereas motor cortex tDCS allowed compensatory changes in both the first and second formant.

A number of studies have recently examined the impact of single-session motor cortex or cerebellar tDCS on visuomotor adaptation. In a seminal study, Galea et al. (2011) provided initial evidence that single-session tDCS has a range of effects on visuomotor adaptation that can be dissociated by changing the location of stimulation. Specifically, tDCS to the cerebellum improved the *speed* of learning, whereas tDCS to motor cortex seemed to increase the *retention* of the learned behaviour after alterations in visual feedback were removed. Improvements in the speed of sensorimotor learning following cerebellar tDCS were also observed in Block and Celnik (2013) and in Doppelmayr et al. (2016). However, in contrast to the results reported by Galea et al. (2011), Doppelmayr et al. (2016) failed to find an effect of motor cortex tDCS on any aspect of sensorimotor learning, including retention. Other work has failed to find differential effects of tDCS on sensorimotor learning related to the stimulated motor area. Herzfeld et al. (2014) found that cerebellar tDCS during sensorimotor learning improved learning speed, but, like Doppelmayr et al., they do not find any impact of motor cortex tDCS on retention. Panouilleres et al. (2015) reported that motor cortex stimulation improved the speed of visuomotor learning, while cerebellar stimulation seemed to have no effect—nearly the opposite pattern of results to that reported by Galea et al. (2011).

By the end of 2016, the weight of published evidence suggested that, more often than not, single-session cerebellar tDCS improved sensorimotor learning, whilst the effects of motor cortex stimulation were no different from sham stimulation. However, Jalali et al. (2017) recently reported that the impact of cerebellar stimulation on sensorimotor learning is highly inconsistent, if present at all. In that paper, the authors use a similar experimental setup to Galea et al. and replicate the finding that cerebellar stimulation speeds up visuomotor learning. But in a series of experiments in which seemingly inconsequential parameters of the task were changed (e.g. the orientation of the screen on which digital representations of limb position are displayed), they found no impact of cerebellar tDCS on sensorimotor learning.

In nearly all the studies cited above, the impact of single-session tDCS on sensorimotor learning was examined by having participants interact with manipulanda to control an abstract representation of limb position (e.g. a cursor on a computer screen). What the detailed experiments of Jalali et al. (2017) suggest is that the nature of this interaction might muddy the impact of cerebellar and motor cortex tDCS on sensorimotor learning. By studying sensorimotor learning in speech we have avoided this confound. Our results are broadly in-line with positive tDCS-related effects noted in limb movement studies and related related work examining the role of the cerebellum in motor control. For instance, stimulating the cerebellum led to improvements in learning that were specific to the correction of sensory prediction errors (Wolpert et al., 1998; Tseng et al., 2007; Galea et al., 2011; Schlerf et al., 2012b; Sokolov et al., 2017). More generally, the results suggest that motor cortex and cerebellar tDCS may have both shared and dissociable effects on sensorimotor learning. This supports the idea that stimulating either structure engages a cortico-cerebellar network involved in the learning and maintenance of motor behaviours.

A post-hoc analysis suggested that, compared to cerebellar tDCS, motor cortex tDCS steered learned changes in speech production closer to previously produced speech utterances. This result needs to be interpreted with caution and tested with further experimentation. Consistent changes in F2 in response to F1 alterations are not omnipresent in the speech motor learning literature (although they may be underreported). For instance, Villacorta et al. (2007) did not observe a change in F2 following an F1 alteration of the same vowel sound used in this study. However, MacDonald et al. (2011) did report F2 changes following F1 alterations. In this case, the authors concluded that the F2 change—which, as in the current study, persisted after altered feedback was removed—was of a different character to changes in F1 that directly compensate for the induced acoustical error. F2 changes in response to F1 alterations were also observed in Rochet-Capellan and Ostry (2011). In this case, during productions of “head”, two different groups of participants experienced alterations of F1 in different directions, and learned changes in F2 were observed in the opposite direction to learned changes in F1.

The pattern of behaviour observed here and in other work is consistent with the idea that compensatory F1-F2 changes in speech related to focal formants manipulations can steer speech towards previously learned speech motor plans. During productions of ‘head’ participants normally decrease their F1 productions in response to an experimentally induced F1 increase. Such a change corrects for the perceived error in the sound of the voice, but it moves speech away from previously learned articulations associated with the F1-F2 vowel space between ‘hid’, ‘head’, and ‘had’. In this case, an increase in F2 concurrent with the decrease in F1 would place articulation closer to a previously learned production (e.g. ‘hid’) in this vowel space. This is precisely what we observed in the group that received motor cortex stimulation, whereas tDCS to the cerebellum seemed to inhibit this behaviour. One interpretation of this result is that precise compensation for F1 alterations versus more global F1-F2 change has a neurological basis. However, there is not enough evidence in the current study to firmly support this, and the idea is in need of further investigation. For instance, we would predict that if the study were repeated with the opposite F1 manipulation a similar pattern of results would be observed: learned changes in F2 would occur in the opposite direction to learned changes in F1 (moving speech towards previously learned motor plans) and motor cortex stimulation would enhance this effect.

Do the results reported here have clinical relevance? In a recent study, Jalali et al. (2017) find that cerebellar tDCS has inconsistent effects on visuomotor adaptation. Over seven experiments, the authors replicate one of the central findings of Galea et al. (2011)—that cerebellar tDCS speeds up sensorimotor learning—but then fail to replicate this result when seemingly inconsequential aspects of the task are changed. They conclude that the use of single-session tDCS in clinical populations, where robust and consistent effects are required, should be questioned. We echo this concern. We did observe a positive impact of anodal tDCS on an aspect of sensorimotor learning in speech compared to sham stimulation. However, despite large group sizes (compared to other work in this area), the effect of tDCS was small. In line with Jalali et al.’s findings, in the case of tDCS-related improvements in sensorimotor learning in the first formant, the effect size (Cohen’s d) averaged across the learning and after-effect phases of the experiment was 0.65. The effect of tDCS on speech motor learning was similar to differences in learning induced via behavioural interventions (Lametti et al., 2014a).

A more substantial problem with applying our results to a clinical setting might be that, in the case of anodal stimulation to the cerebellum, we observed both facilitation and inhibition of learning. Despite claims to the contrary, anodal stimulation can impair certain behaviours (and cathodal stimulation can improve them) (Ferrucci et al., 2008; Pope and Miall, 2012; Bestmann et al., 2015; Lametti et al., 2016; Oldrati and Schutter, 2017). Indeed, our understanding of the mechanisms by which tDCS influences behaviour remains largely speculative, and the stimulation-action relationship is likely more complex than is often assumed. One issue is that, due to the spatial resolution at which tDCS operates, stimulation effects are likely not constrained to the target areas and instead engage a network of regions. Furthermore, anodal tDCS may modulate pyramidal and Purkinje cells of the motor cortex and cerebellum respectively, but it may also act on inhibitory interneurons. This could affect behaviour in unexpected ways as the relationship between excitation and inhibition in the affected network is rebalanced (Bestmann et al., 2015). Here we demonstrate that anodal tDCS facilitates certain aspects of a behaviour while inhibiting others, an effect that has been noted before (albeit in a different behavioural context) (Luculano and Cohen Kadosh, 2013). The small effects observed here, and the task-dependent variability in the direction of these effects, make tDCS less than ideal for clinical use, at least in the case of cerebellar stimulation. Thus, our study supports the idea that tDCS is a useful tool for investigating brain-behaviour relationships, but the results provide only weak support for the clinical use of single-session tDCS in speech.

Speech adaptation in this study was notably smaller than that reported in our previous work (e.g. Lametti et al. 2014). This was especially apparent in the case of compensation in the first formant. There are two possible reasons for this. In our previous work a single word was produced during altered feedback. Here participants produced three words during altered feedback. Speech adaptation related to real-time alterations in vowel sounds is partly dependent on the consonants that surround the vowel and transfer of learning from a word like ‘head’ to a word like ‘bed’ is weak (see Rochet-Capellan and Ostry 2010). Having participants produce multiple words slows learning as multiple acoustic targets must be met. F1 adaptation may have also been curtailed by the attachment of tDCS to the head. To replicate previous work, during cerebellar stimulation, the reference electrode was placed on the cheek (Galea et al. 2011). To allow for this, in all stimulation conditions, a flexible self-adherent bandage ran from the bottom of the jaw around the head (in front of the vertex) and back to the bottom of the jaw. The first formant is largely determined by tongue height and changes in tongue height may have been restricted because of this tDCS setup. Future work involving cerebellar tDCS and speech production should consider placing the reference electrode on the shoulder (Panouillères et al., 2015).

In summary, we investigated the impact of motor cortex or cerebellar tDCS on sensorimotor learning in speech. We found that motor cortex and cerebellar tDCS have both shared and dissociable effects on this type of sensorimotor adaptation. Cerebellar tDCS focused sensorimotor adaptation in speech on the source of the acoustical error, whereas motor cortex tDCS allowed more general compensatory changes in speech production. We posit that motor cortex tDCS drives sensorimotor learning towards previously learned patterns of movement and suggest that this idea should be tested further.

## Acknowledgements

This study was funded by postdoctoral fellowships to DRL from the British Academy and Corpus Christi College Oxford and a Medical Research Council UK project grant to KEW (MR/M025539/1).

## Author Contributions

DRL and KEW designed the experiment. HJS and PF recruited, screened, and tested the participants. DRL and HJS analyzed the data, created the figures, and drafted the manuscript. DRL, HJS and KEW edited the manuscript for publication.

